# Mitochondrial Dysfunction is Part of a Novel Mechanism for Chemotherapy-Induced Neuropathic Pain

**DOI:** 10.64898/2026.11.06.764717

**Authors:** Reem Sakas, Saher Abu-Ata, Yara Nakhleh-Francis, Nasser Abd Elkarim, Akhilesh Kotiyal, Henry Politte, Yaseen Awad-Igbaria, Jean F. Soustiel, Eilam Palzur

**Affiliations:** Azrieli Faculty of Medicine, Bar-Ilan University, Zefat, Israel; Research Institute of Galilee Medical Center, Nahariya, Israel; Department of Obstetrics and Gynecology, Galilee Medical Center, Nahariya, Israel Haifa, Israel; General Intensive Care Department at Hadassah Ein-Kerem, Jerusalem, Israel; Department of Anesthesiology, Center for Clinical Pharmacology, Washington University in St. Louis, St. Louis, MO, USA; Department of Chemistry, School of Science and Engineering, Saint Louis University, St. Louis, MO, USA; Department of Neurosurgery, Galilee Medical Center, Nahariya, Israel

**Keywords:** Paclitaxel-induced peripheral neuropathy, Neuropathic pain, Mitochondrial function, TRPV1, Spinal cord, mPTP

## Abstract

Chemotherapy-induced peripheral neuropathy (CIPN) is a common, persistent complication of cancer treatment, yet the mechanisms sustaining pain remain unclear. Mitochondrial dysfunction has been implicated in chronic pain, but its role in spinal cord dysfunction after chemotherapy, and whether it can be reversed, is unknown. Here, we show that paclitaxel-induced persistent mechanical allodynia was associated with spinal cord (L1-L6) mitochondrial dysfunction 18 days after the last injection, including reduced oxygen consumption and membrane potential (Ψm) in the non-phosphorylated state, ATP-linked respiration, maximal respiration, and electron transfer capacity. Paclitaxel also increased susceptibility to mitochondrial permeability transition pore (mPTP)-induction, lowering calcium retention capacity at 3, 8, and 13 days after the last injection. Acute ex-vivo paclitaxel exposure had a similar effect, and cyclosporin A, an mPTP blocker, significantly reversed it. Consistent with a causal role for mitochondrial impairment, hyperbaric oxygen therapy (HBOT), initiated after allodynia developed, improved oxygen consumption and Ψm and progressively enhanced mechanical thresholds. Likewise, intrathecal delivery of healthy mitochondria to the spinal cord attenuated established allodynia in paclitaxel-treated rats and reduced mechanical and thermal hypersensitivity in other pain models, sciatic nerve crush and capsaicin. These findings identify persistent spinal mitochondrial dysfunction and increased mPTP susceptibility as features of CIPN and support mitochondrial protection as a therapeutic strategy for CIPN and other pain conditions.

## Introduction

Mitochondria play a critical role in maintaining neuronal function and homeostasis by ensuring health metabolic functions and energy production via oxidative phosphorylation (OXPHOS) process^1^. Moreover, vital neuronal processes such as calcium homeostasis, ion channel activity, and reactive oxygen species (ROS) signaling are regulated by mitochondria^2^. Deficits in mitochondrial function have been suggested as a potential factor contributing to the development of chronic pain^3,4^. This emerging perspective has paved the way for novel pharmacological and non-pharmacological therapeutic strategies targeting mitochondrial dysfunction^4–8^. The central hypothesis underlying these strategies is that restoring and protecting mitochondrial function can prevent the transition from acute to chronic pain by re-establishing neuronal homeostasis in sensory pathways^9^. However, it remains unclear whether improving mitochondrial function after pain has developed can still produce analgesic effects.

Previous evidence suggests chemotherapy such as paclitaxel induces mitochondrial dysfunction in the sensory pathways, contributing to the development of chronic pain^10,11^. One proposed mechanism is that paclitaxel interacts directly with mitochondria and can promote the mitochondrial permeability transition pore (mPTP) induction. This may dissipate the mitochondrial membrane potential, impair Ca² retention and oxidative phosphorylation, reduce ATP production, and increase ROS generation^10,12^. Yet, this proposed mechanism remains under investigation and has not been fully validated. Accordingly, we sought to determine whether and how paclitaxel induces mitochondrial dysfunction in the spinal cord, and whether restoring mitochondrial function can attenuate peripheral allodynia across three established preclinical pain models: paclitaxel-induced neuropathy, sciatic nerve crush, and capsaicin-induced hypersensitivity.

## Materials and Methods

### Animals

Animal study was approved by the Animal Care Committee of Bar-Ilan University. Male Sprague-Dawley rats (200-280g; 7-8 weeks age) were used in this study. Animals were housed in groups of 4 in a sterilized solid bottom cage with contact bedding under controlled temperature and a 12:12h light/dark cycle.

### Chemotherapy induced peripheral neuropathy (CIPN)

Paclitaxel solution (6mg/ml) was dissolved with physiological saline and administered every other day for 7 days (i.e., days 1, 3, 5, 7) at a dose of 2mg/kg (IP).

### Neuropathic pain model

Rats were anesthetized using isoflurane (2.5% in 00% oxygen). The sciatic nerve was exposed by performing a 1.5 cm incision in the skin on the posterior face of the thigh, creating a gap in the thigh muscle. The exposed nerve was crushed by applying pressure with micro-dissecting tweezers for 15 seconds^13^. Afterward, the muscle and skin were then sutured, and rat was placed in an individual cage and monitored until recovery from anesthesia.

### Mitochondria isolation

Brain or spinal cord was extracted (L1-L6) and washed twice with respiration buffer MiR05^14^. Then, the tissue samples were cut and homogenized with cold extraction buffer-A, and centrifuged at 600g for 10 min at 4 °C. Then, the supernatant was collected and re-centrifuged at 11,000g for 10 min at 4 °C. Pellets were resuspended with a second cold extraction buffer-B and processed by sequential centrifugation as above. Finally, the pellet was then resuspended in saline for transplantation or MiR05 for mitochondrial function assessment, and the mitochondrial concertation was determined according to the Bradford assay using the Pierce, Coomassie Plus Protein Assay with BSA as a standard.

### Drug preparation and intrathecal injection

Capsaicin and AMG-9810 were dissolved in DMSO (1% of the final volume), and the stock solution was dissolved in saline. Under brief anesthesia (isoflurane 1.5%), capsaicin (50µg/30µl), AMG-9810 (50µg/30µl), or isolated mitochondria (100µg/30µl) were administered intrathecally (i.t) via lumbar puncture between the L4 and L5 vertebrae^14^.

### HBO treatment

The Hyperbaric oxygen chamber (Hipertech Inc, Istanbul, Turkey) was flushed with 100% oxygen before the animals were placed inside. The chamber pressure was then increased to 2.5 atmospheres absolute (ATA) at a rate of 0.1 ATA/min and maintained at this pressure for 90 minutes.

### Behavioral tests and pain assessment

Behavioral testing was conducted under red light illumination during the dark phase of the cycle. Animals were habituated to the testing room for 1h before testing. All behavioral assessments were performed by an experimenter blinded to the experimental conditions.

#### Mechanical sensitivity

The mechanical threshold of the right hind paw was measured using an electronic von Frey apparatus (cat. No. 38450; Ugo Basile). Paw lifting or shaking in response to stimulation was considered a positive response. Five measurements were collected from each animal.

#### Thermal sensitivity

Thermal sensitivity was assessed using a hot-plate analgesia meter (cat. No.3515-022; Ugo Basile). Rats were acclimated to the chamber at 35°C for 10min, after which the plate temperature increased at a rate of 1°C/min. Thermal responses, including hind-paw licking, jumping, and rearing, were recorded.

#### Open field

The animals were placed in one corner of the open field arena (L70 × W70 × H50 cm). The session lasted 20 minutes, and locomotor activity measurements and time spent in the zone were recorded and measured automatically using EthoVision (V.15).

#### Motor outcome

Motor coordination was assessed using the rotarod apparatus (Model 47750; Ugo Basile). Animals were trained and tested with rotarod before the intervention. In the testing trials, the rotation speed was gradually increased from 4 to 40 rpm over 300 seconds. Each trial ended when the animal fell from or clung to the rod, or when the maximum duration of 300 seconds was reached^13^.

### Mitochondrial function

#### Respiration measurements and mitochondrial membrane potential-Ψm

High-resolution respirometry was performed on fresh tissue using the OROBOROS-O2k (Oroboros Instruments, Innsbruck, Austria). To evaluate mitochondrial membrane potential (Ψm), safranin calibration was performed before the experiments. Mitochondrial oxygen consumption and Ψm were evaluated in routine state, non-phosphorylation state (Leak), phosphorylation state (OXPHOS), and electron transfer (ET) capacity (Detailed methods are reported in the Supplementary material).

#### Ca^2+^ retention capacity (CRC) in spinal cord tissue

The spinal cord (L1-L6) was harvested, weighed, and washed with Tris-HCl buffer at 4 °C. Afterward the tissue samples were homogenized using a pestle and glass with ice-cold Tris-HCl buffer. The CRC was examined using the O2k-CaGreen-5N module, in the presence of complex I inhibitor, complex II substrate, and ADP. 50µL of tissue homogenate was added, followed with 20µm of CaCl_2_, and then we used stepwise CaCl_2_ titration of 10µm, approximately every 4-5 min. Ca^2+^ uptake was considered when fluorescence signals exhibited an exponential-like decay shape. The induction of mPTP was identified by an increase in fluorescence signals without added Ca^2+^ boluses or by stabilizing the fluorescence signal after Ca^2+^ titration (Detailed methods, substrate/inhibitor concentration and protocol are reported in the Supplementary material).

### Data analysis

Statistical analyses were performed using GraphPad Prism (v10). Differences between two groups were assessed using Unpaired t-tests. The effect of condition and different treatments were assessed using two-way ANOVA. The effect of conditions, different treatments, and different time-points were assessed using two or three-way ANOVA. Significant main effects and interactions were further pursued using Holm-Sidak’s post hoc test. The accepted significance value for all tests was set at *P*<0.05. All data were expressed as Mean ± SEM.

## Results

### HBOT reverse peripheral allodynia following paclitaxel treatment

Here, we examined whether HBOT can reverse peripheral allodynia induced by paclitaxel. Rats were injected with paclitaxel (2mg/kg IP) every other day for 7 days (Fig. 1A), HBOT was performed twice a day from day 9 to 25 (Fig. 1A), and mechanical threshold was measured on day 9, 12, 15, 20 and 25, before the first HBOT session (Fig. 1A). Before the HBOT, the paclitaxel-vehicle and paclitaxel-HBOT group showed a significant reduction in the hind paw mechanical threshold compared to the baseline, with no significant difference between the groups (Fig. 1B). Following 3 days of HBOT, we observed a significant improvement in mechanical threshold of the paclitaxel-HBOT group (Fig. 1B). By day 15, corresponding to 6 days of HBOT, the mechanical threshold had increased further and remained stable through days 20 and 25 (Fig. 1B). Thus, HBOT progressively reversed paclitaxel-induced mechanical allodynia, and this improvement was sustained after treatment. Notably, reduction in motor performance was observed only in the paclitaxel-vehicle group on day 25 compared to the baseline (Fig. 1C), with no difference between the groups in the exploration and locomotion test (Fig. 1D).

**Figure 1.**
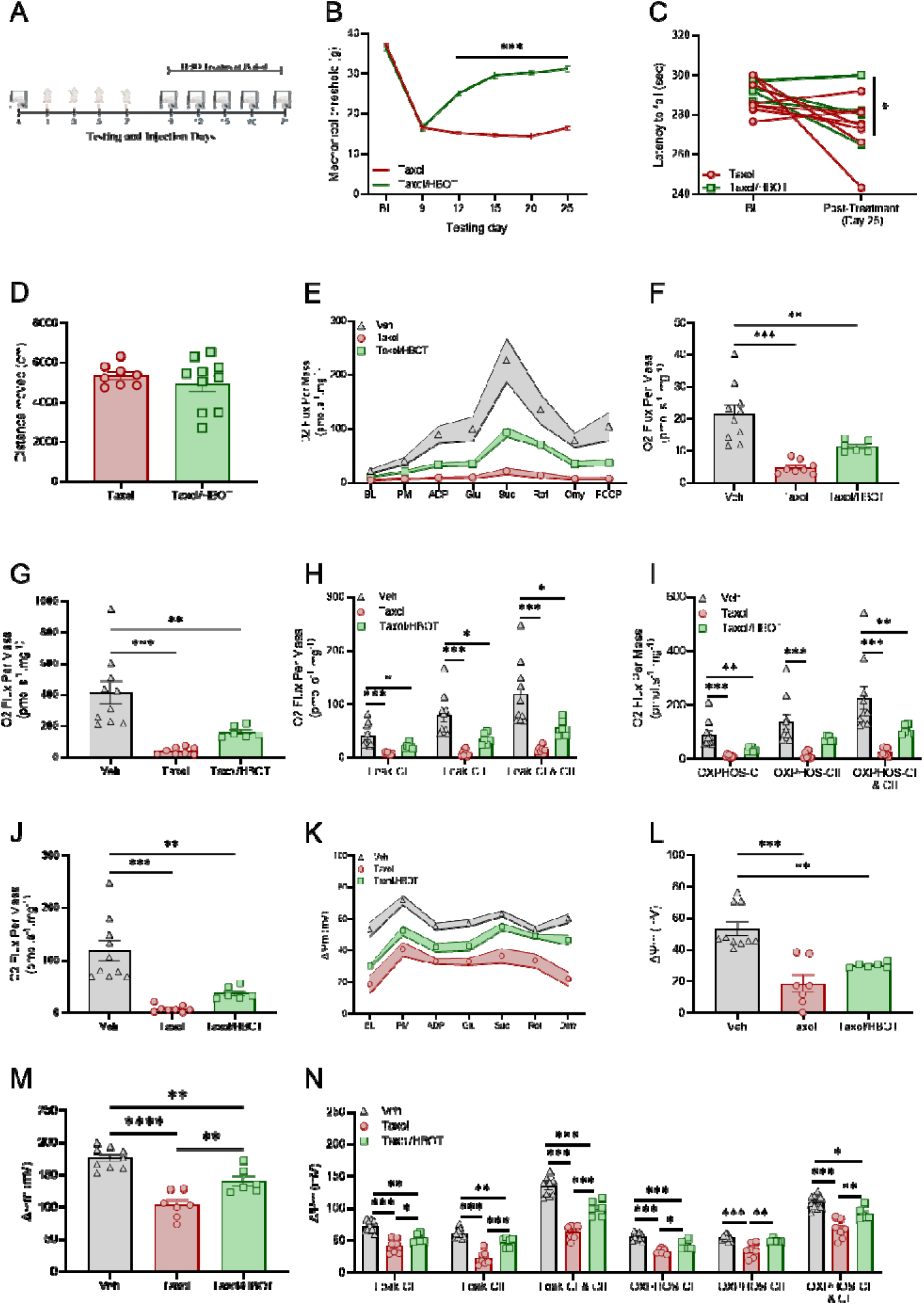
HBO treatment reverses peripheral allodynia in CIPN model via protecting of the mitochondrial function. **(A)** The experimental timeline. Mechanical and thermal sensitivity, and motor performance were measured before paclitaxel. Animals were injected with paclitaxel (2mg/kg, IP) every other day for 7 days (i.e., days 1, 3, 5, 7). On day 9, following hind paw mechanical threshold assessment, animals were randomly assigned to one of the experiment groups: paclitaxel group, paclitaxel/HBOT group. The HBO treatment was performed twice a day from day 10 to 25. The mechanical sensitivity was measured before the HBOT session on day 9, 12, 15, 20 and 25. Motor performance, and locomotion were assessed on day 25-26. (**B**) Mechanical force withdrawal (G). **(C)** Motor performance in rotarod test expressed by latency to fall (Second). **(D)** Total distance moved in the OF test (cm). **(E)** Representative respirometric traces from spinal cord mitochondria using a substrate-uncoupler-inhibition protocol to measure oxygen consumption and Ψm on day 25 in the paclitaxel-induced CIPN model. **(F)** Routine mitochondrial respiration (without stimulators or inhibitors). **(G)** Maximal mitochondrial respiration (CI+CII). **(H)** Mitochondrial respiration in the OXPHOS state coupled with phosphorylation of ADP to ATP (CI, CII, CI+CII). **(I)** Mitochondrial respiration in the non-phosphorylation state (Leak). **(J)** Mitochondrial respiration in the Electron transfer (ET) capacity state. **(K)** Representative Ψm mean trace from spinal cord mitochondria using a substrate-uncoupler-inhibition protocol to examine mitochondrial membrane potential by safranin. **(L**) Basal Ψm (without stimulators or inhibitors). **(M)** The Ψm in maximal mitochondrial respiration (CI+CII). **(N)** The Ψm in the non-phosphorylation state (Leak), and the OXPHOS state coupled with phosphorylation of ADP to ATP. Mixed Model ANOVA Followed with Holm-Sidak’s test, One-way ANOVA followed with Holm-Sidak’s test, unpaired t-test. Mean ± SEM. \**P<*0.05, \*\**P<*0.01, \*\*\**P<*0.001, Difference between groups.

### Paclitaxel-induced spinal cord mitochondrial dysfunction

We examined mitochondrial function in terms of oxygen consumption and Ψm on day 26 (18 days after the last paclitaxel treatment on day 7, Fig. 1A). We observed a significant reduction in mitochondrial oxygen consumption and Ψm in the paclitaxel-vehicle group at the basal (Fig. 1E-F, K-L), maximal respiration (Fig. 1G, M), non-phosphorylation (Fig. 1H, N), phosphorylation (Fig. 1I, N), and electron transfer (ET) capacity state (Fig. 1J) compared to the vehicle group (Fig. 1E-N). HBOT has shown a significant improvement in the Ψm (Fig. 1E-N) and tended on average to show higher mitochondrial oxygen consumption compared to the paclitaxel-vehicle group (Fig. 1E-N).

### Paclitaxel facilitates mPTP induction

Paclitaxel can lead to a robust Ca^2+^ influx in cells. We speculate that paclitaxel enhances the intracellular Ca² levels, leading to mitochondrial Ca² overload, Ψm depolarization, and subsequent induction of the mPTP. Consequently, we measured the mitochondrial Ca^2+^ retention capacity (CRC) in the spinal cord (L1-L6) isolated from animals that were treated with vehicle or paclitaxel (2mg/kg, IP) every other day for 7 days.

On day 10, 15, 20, CRC was measured in spinal cord tissue homogenates with/without the presence of 2µm of Cyclosporin A (CsA, a known mPTP inhibitor, Fig. 2A). A significant impairment in the CRC was observed following paclitaxel treatment on day 10, 15, and 20 compared to Vehicle treatment (Fig. 2B-F). In the vehicle group (in the absence of CsA), the mPTP-induction was observed after adding 82±5.83 µm of Ca^2+^ (Fig. 2B, F). As excepted CsA (2 μM) delayed mPTP induction, increasing the cumulative Ca² load to 100 ± 3.16 μM (Fig. 2B, F). However, paclitaxel treatment leads to lower CRC at the three time-points (day 10, 15, and 20, Fig. 2C-F). mPTP induction (in the absence of CsA) was observed following adding 52.5±7.5 µm of Ca^2+^ (on day 10, Fig. 2C, F), 47.5±4.78 µm of Ca^2+^ (on day 15, Fig. 2D, F), and 50±5.77 µm of Ca^2+^ (on day 20, 2E, F). CsA treatment significantly delayed mPTP induction, with no significant interaction (Fig. 2B-F). Similar early mPTP induction was observed after a 20-min incubation of spinal cord tissue with 20µM paclitaxel ex-vivo (Fig. 2G-I). Together, these findings indicate that paclitaxel facilitates mPTP opening in the spinal cord, likely contributing to neuronal damage.

**Figure 2.**
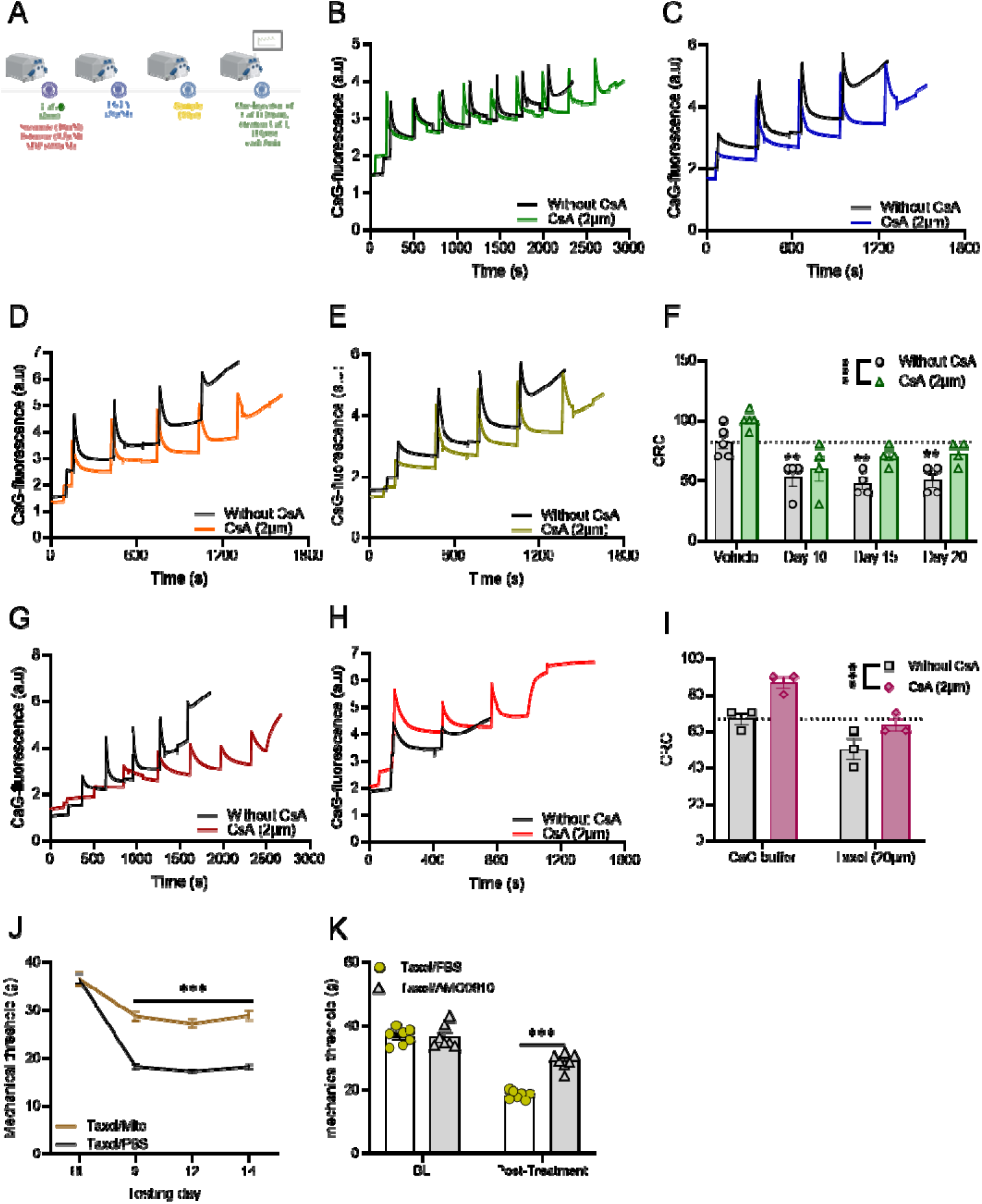
Paclitaxel facilitates mPTP induction, mitochondrial transplantation and blocking TRPV1 in the spinal cord attenuate mechanical allodynia. **(A)** The experimental timeline of CRC. Following the addition of CaG (2 μM), succinate (10 mM), ADP (600 μM), rotenone (0.5 μM), EGTA (50 μM), 50 μl of the spinal cord tissue homogenate was added to the chamber, followed with a single injection of CaCl2 (20 μM), and a titration of CaCl2 (10 μM). (B, C, D, E) Representative Ca²-uptake traces from naïve rats (B) and paclitaxel-treated rats on days 10 **(C)**, 15 **(D)**, and 20 **(E)**, measured without or with CsA (2μM). Each peak represents a CaCl addition; the subsequent decrease in CaG fluorescence reflects mitochondrial Ca² uptake. **(F)** Quantification of CRC, defined as the cumulative Ca² load tolerated before mPTP induction, for the corresponding groups. CsA significantly increased CRC, paclitaxel-treated mice showed a significant reduction in CRC at days 10, 15 and 20 compared with the vehicle (without CsA) group. **(G, H)** Representative Ca²-uptake traces following spinal cord tissue incubation with CaG buffer **(G)** or paclitaxel (20µM) for 20minuites **(H)**, measured without or with CsA (2μM). **(I)** Quantification of CRC, defined as the cumulative Ca² load tolerated before mPTP induction, following spinal cord tissue incubation. **(J)** Hind paw mechanical thresholds measured at baseline (BL) and on days 9, 12, and 14 in rats treated with paclitaxel (2 mg/kg, IP) every other day for 7 days. On day 9 healthy mitochondria (100μg/30 μL; Taxol/Mito) or vehicle (30 μL; Taxol/PBS) intrathecally on day 8. **(K)** Baseline and post-treatment thresholds in paclitaxel-treated rats given the TRPV1 antagonist AMG9810 or PBS intrathecally on day 9. Mixed Model ANOVA followed with Holm-Sidak’s test, Two-way ANOVA followed with Holm-Sidak’s test. Mean ± SEM. \**P<*0.05, \*\**P<*0.01, \*\*\**P<*0.001, Difference between groups. \*\**P<*0.01 compared to the vehicle/without CsA group.

### Transplantation of healthy mitochondria into the spinal cord reverses peripheral allodynia

Although spinal TRPV1 signaling has been shown to mediate mitochondrial dysfunction, and increased TRPV1 expression has been reported in the spinal cord following paclitaxel treatment, it remains unclear whether restoring spinal mitochondrial bioenergetics can alleviate paclitaxel-induced mechanical hypersensitivity. To test whether mitochondrial transplantation could reduce established hypersensitivity, we measured baseline hind-paw mechanical thresholds and then treated rats with paclitaxel every other day for 7 days. On day 8, freshly isolated naïve-rat brain mitochondria (100 µg/30 µL) or vehicle (30 µL) were administered intrathecally by lumbar puncture (L4-L5) vertebrae. Hind paw mechanical thresholds were reassessed on days 9, 12, and 14. The results demonstrated that mitochondrial transplantation attenuated paclitaxel-induced mechanical allodynia, as rats in the paclitaxel-mitochondria group exhibited significantly higher mechanical thresholds on days 9, 12, and 14 compared to rats in the paclitaxel-vehicle group (Fig. 2J).

Consistent with a role for TRPV1 signaling, intrathecal administration of the TRPV1 antagonist AMG9810 at L4–L5 on day 9 significantly increased hind-paw mechanical thresholds in rats treated with paclitaxel every other day for 7 days, compared with Taxol/PBS group, indicating attenuation of paclitaxel-induced mechanical allodynia (Fig. 2K). Together, these findings support spinal mitochondrial dysfunction and TRPV1 signaling as potential therapeutic targets for chemotherapy-induced neuropathic pain.

To determine whether the beneficial effect of mitochondrial transplantation extended beyond CIPN, we tested it in two additional pain models: intrathecal capsaicin administration and SNC, that we previously showed to produce allodynia accompanied by mitochondrial dysfunction in the spinal cord^6,13,14^.

Following SNC, there was a significant reduction in mechanical threshold, that was observed on day 2 following nerve injury (Fig. 3A-B). On day 3, fresh mitochondria (100µg/30 µL) or vehicle (30µL) were administered intrathecally by L4 and L5 vertebrae (Fig. 3A). Hind paw mechanical thresholds were assessed on days 5 and 7, thermal hypersensitivity and motor function were evaluated on day 8 (Fig. 3A-E). Mitochondrial transplantation reversed mechanical allodynia, as rats in the SNC-mitochondria group exhibited significantly higher mechanical thresholds on days 5 and 7 compared to rats in the SNC-vehicle group (Fig. 3B). The SNC-mitochondria group also exhibited reduced thermal hypersensitivity (Fig. 3C) and improved motor performance compared with the SNC-vehicle group (Fig. 3D). However, locomotor activity did not differ between the groups following mitochondrial or vehicle administration (Fig. 3E). A similar pattern was observed in the capsaicin model, in which mitochondrial transplantation increased mechanical threshold on days 5 and 7 and reduced thermal hypersensitivity on day 8 compared with vehicle treatment (Fig. 3F-I). Notably, in both models, we confirmed impaired spinal cord mitochondrial respiration and altered Ψm before transplantation on day 2 (Fig. 4A-J).

**Figure 3.**
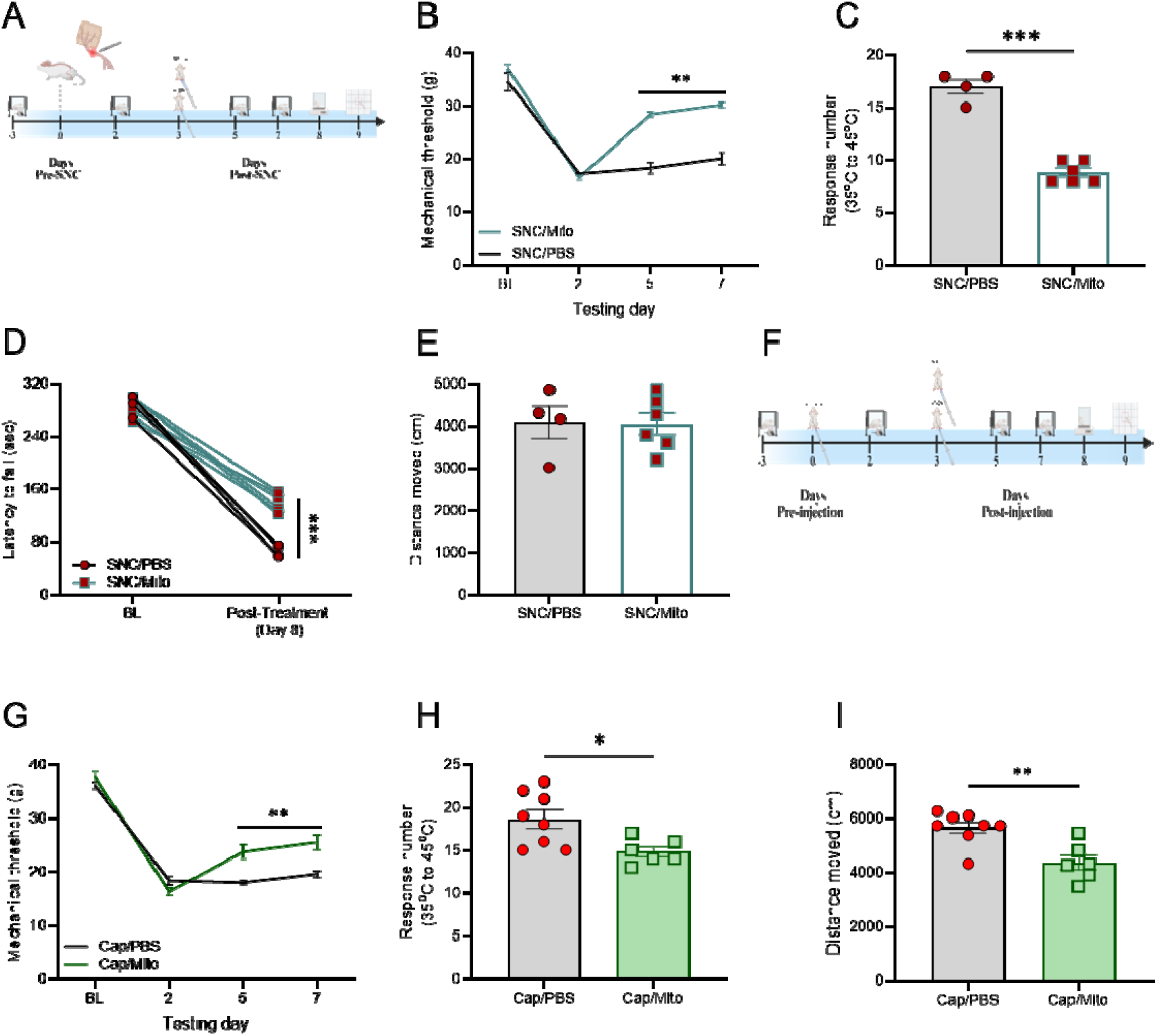
Healthy mitochondrial transplantation into the spinal cord reverses peripheral hypersensitivity in pain models. **(A)** The experimental timeline. Mechanical sensitivity was measured Pre- and Post-SNC on day 2. 3 days following SNC, mitochondria (100ug in 30ul) were injected into the spinal cord. Mechanical sensitivity was measured on day 5, and 7. Thermal sensitivity was measured on day 8. (**B**) Mechanical force withdrawal (G). **(C)** Sum of nociceptive response on day 8 post SNC. **(D)** Motor performance in rotarod test expressed by latency to fall (Second). **(E)** Total distance moved in the OF test (cm). **(F)** The experimental timeline. Mechanical sensitivity was measured Pre- and Post-Cap injections on day 2. Following Cap injection on day 3, mitochondria (100ug in 30ul) were injected into the spinal cord. Mechanical sensitivity was measured on day 5, and 7. Thermal sensitivity was measured on day 8. (**G**) Mechanical force withdrawal (G). **(H)** Sum of nociceptive response on day 8 post Cap injection. **(I)** Total distance moved in the OF test (cm). Mixed Model ANOVA followed with Holm-Sidak’s test, unpaired t-test. Mean ± SEM. \**P<*0.05, \*\**P<*0.01, \*\*\**P<*0.001, Difference between groups.

**Figure 4.**
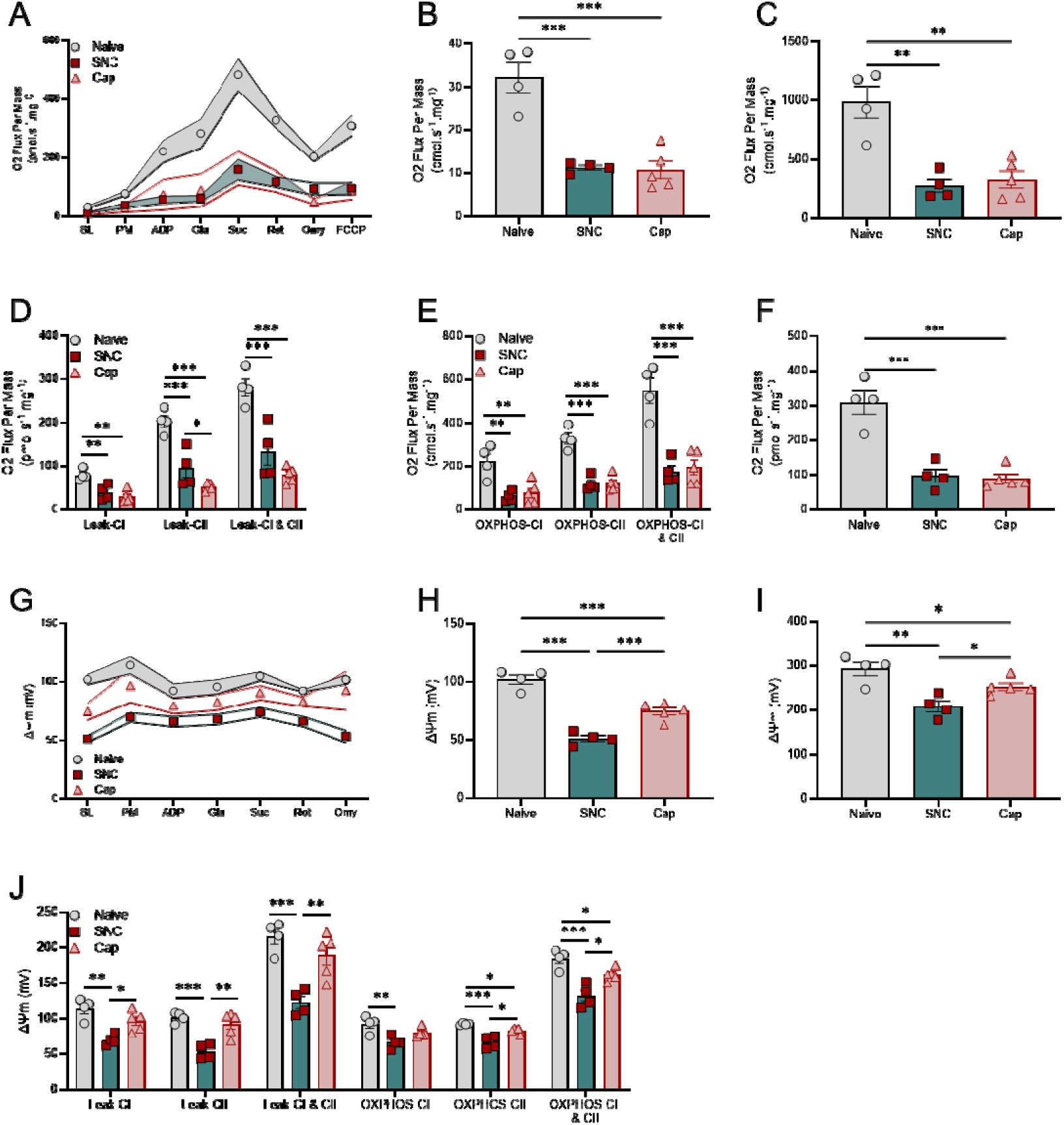
Mitochondrial oxygen consumption and Ψm in the spinal cord 2 days following SNC, and capsaicin injection. **(A)** Representative respirometric traces from spinal cord mitochondria using a substrate-uncoupler-inhibition protocol to measure oxygen consumption and Ψm. **(B)** Routine mitochondrial respiration (without stimulators or inhibitors). **(C)** Maximal mitochondrial respiration (CI+CII). **(D)** Mitochondrial respiration in the non-phosphorylation state (Leak). **(E)** Mitochondrial respiration in the OXPHOS state coupled with phosphorylation of ADP to ATP (CI, CII, CI+CII). **(F)** Mitochondrial respiration in the Electron transfer (ET) capacity state. **(G)** Representative Ψm mean trace from spinal cord mitochondria using a substrate-uncoupler-inhibition protocol to examine mitochondrial membrane potential by safranin. **(H**) Basal Ψm (without stimulators or inhibitors). **(I)** The Ψm in maximal mitochondrial respiration (CI+CII). **(J)** The Ψm in the non-phosphorylation state (Leak), and the OXPHOS state coupled with phosphorylation of ADP to ATP. One-way Anova followed with Holm-Sidak’s test. Mean ± SEM. \**P<*0.05, \*\**P<*0.01, \*\*\**P<*0.001.

## Discussion

Paclitaxel produced persistent mechanical hypersensitivity, reduced mitochondrial oxygen consumption and Ψm, and lowered the mitochondrial Ca² uptake of the spinal cord. Conversely, preserving mitochondrial function with HBOT reversed mechanical allodynia. We further showed that directly supporting neuronal health by delivering healthy mitochondria to the spinal cord also reversed paclitaxel-induced allodynia, and this intervention attenuated allodynia in two mechanistically distinct pain models

Paclitaxel can injure and degenerate sensory axons, leading to the development of peripheral hypersensitivity^15^, which can be attenuated by concurrently with HBOT^16^. We found that HBOT significantly reversed allodynia in the CIPN model, an effect that believed to be depend on downregulating spinal TRPV1 expression^16^. The upregulation of spinal TRPV1 expression and signaling in chronic pain conditions including CIPN, is crucial pathway that leads to spinal mitochondrial dysfunction-mediated chronic pain development, a mechanism that previously confirmed^14^, as we expected blocking spinal TRPV1, which protect mitochondria function in chronic pain^14^, reversed paclitaxel-induced allodynia.

We found a reduction in oxygen consumption and Ψm at basal, maximal respiration, phosphorylation, non-phosphorylation respiration, and ETC, measured 18 days after the last paclitaxel injection. The widespread damage in mitochondrial function suggests a generalized loss of mitochondrial capacity rather than a defect confined to a specific respiratory complex or state. The persistence of these effects after paclitaxel treatment indicates a sustained bioenergetic deficit in the spinal cord. This deficit could contribute to central sensitization because neurons have high energy demands and rely on oxidative phosphorylation to support synaptic transmission, and maintain ionic homeostasis^17^. HBOT improved mitochondrial oxygen consumption and Ψm after paclitaxel treatment, suggesting that this dysfunction is amenable to intervention, this effect may resulting from activation antioxidant defenses pathways and mitochondrial biogenesis^18^.

Paclitaxel is known to increases swollen and vacuolated mitochondria in sensory axons^15^, impairs respiration and ATP production in peripheral nerve^19^, reduces respiratory capacity in sensory pathway^19–22^. These effects may reflect disrupted Ca² homeostasis, as paclitaxel promotes Ca² influx and release from intracellular^23,24^, leading to mitochondrial Ca² overload, which facilities mPTP induction^10^, resulting in collapses Ψm, depletes ATP, and causes mitochondrial swelling^15,25,26^. Paclitaxel may also act directly on mitochondria, and trigger mitochondrial Ca² release through mPTP induction^12^, as it interacts with mitochondrial-β-tubulin associated with VDAC in the mPTP complex^27^, and induces cytochrome-c release from mitochondria, an effect found to be blocked by the mPTP inhibitor-CsA^27,28^. Supporting the involvement of mPTP and a direct action of paclitaxel on mitochondrial membranes, independent of its canonical effects on microtubules. Here, we found that paclitaxel markedly reduced mitochondrial respiration and Ψm, and early mPTP induction long after the last injection. And CsA treatment significantly delays mPTP induction. Notably, incubating spinal cord tissue with paclitaxel reproduced the increased susceptibility to mPTP induction. Together, these findings provide convergent evidence that paclitaxel induces long-lasting mitochondrial dysfunction, reflected in part by increased susceptibility to mPTP induction. The effect of paclitaxel in spinal cord tissue suggests that it may act directly on spinal cord mitochondria.

Improving mitochondrial function to support neuronal health are emerging as potential therapeutic strategies for spinal cord injury, and traumatic brain injury (TBI)^29,30^. Here, we found that intrathecal delivery of healthy mitochondria after the development of peripheral hypersensitivity-induced by paclitaxel significantly increased mechanical thresholds. This intervention has also been shown to attenuate allodynia in other pain models associated with spinal cord mitochondrial dysfunction. These findings are consistent with evidence that extracellular mitochondria can be taken up by recipient cells and may help restore bioenergetic function, buffer intracellular Ca², and reduce oxidative stress and inflammatory signaling, this intrinsic mechanism has been found to be involved in pain resolution^4,9^.

Our findings identify spinal cord mitochondrial dysfunction, characterized by impaired respiration, depolarization, and heightened susceptibility to mPTP induction, as a consequence of paclitaxel treatment and a contributor to CIPN. Crucially, restoring mitochondrial function reverses established mechanical allodynia in different pain models. These results support mitochondrial protection as a therapeutic strategy for CIPN and for chronic pain conditions that involve mitochondrial impairment.

## Declarations

### Ethics approval

Animal study was approved by the Bar-Ilan University Animal Care Committee and was carried out in accordance with the National Institutes of Health Guide for the Care and Use of Laboratory Animals.

### Consent for publication

Not applicable.

### Availability of data and materials

The datasets included in the study, and the code for statistical analysis are available from the corresponding author upon request.

### Conflict of interest statement

The authors declare no competing interests.

### Funding

This study was supported by the Excellence Research Award granted to RS.

### Author contributions

Conceptualization YAI., RS; Methodology RS., SAA., AK., HP., YNF., NAE., EP; Investigation YAI., RS., SAA; Visualization YAI., JFS., EP; Funding acquisition: RS; Project administration YAI; Writing – original draft YAI., RS; Writing – review & editing YAI., JFS., EP.

## Acknowledgments

Ms. RS as well, all authors would like to dedicate this work to the memory of Suhela Sakas whose strength, kindness, and unwavering support continue to inspire the team. Though she is no longer with us, yet her spirit remains a guiding light in all that we do.

## Abbreviations

CIPN: Chemotherapy-induced peripheral neuropathy
CRC: Ca² retention capacity
CsA: Cyclosporin A
DRG: Dorsal root ganglion
ET: Electron transfer
HBOT: Hyperbaric oxygen therapy
mPTP: Mitochondrial permeability transition pore
OXPHOS: Oxidative phosphorylation
ROS: Reactive oxygen species
SNC: Sciatic nerve crush
TBI: Traumatic brain injury
TRPV1: Transient receptor potential vanilloid-1
VDAC: Voltage-dependent anion channel
Ψm: Mitochondrial membrane potential

## Notes

### Competing Interest Statement

The authors have declared no competing interest.

